# Exploiting Index Cross-Talk to Modify Variant Calls

**DOI:** 10.1101/332346

**Authors:** Peter M. Ney, Lee Organick, Karl Koscher, Tadayoshi Kohno, Luis Ceze

## Abstract

Modern next-generation DNA sequencers support multiplex sequencing to improve throughput and decrease costs. This is done by pooling and sequencing samples together in parallel, which are later demultiplexed according to their unique indexes^1, 2^. When reads are assigned to the wrong index, called index cross-talk, information is leaked between samples^3–6^. This creates a physical information side-channel, a well known class of vulnerabilities in information security^7–10^, that may be used to modify downstream results. Here we demonstrate the feasibility of such an attack through the use of a separately indexed library that causes a wild-type human exome to be misclassified as heterozygous at the sickle-cell locus. Simple methods can be used to minimize or detect attempts to modify genetic variants using this side-channel, such as filtering by read quality or finding outliers in read coverage. To further minimize this risk we recommend the use of new library preparation methods that reduce index cross-talk, like unique dual indexes^11, 12^, whenever samples are sequenced together in important applications. Biotechnology that interfaces molecular and digital information, like DNA sequencers, may have security risks typically associated with information systems, including the side-channel vulnerability described in this study. We encourage the community to consider the security of genomics-information pipelines before they reach mass adoption.

---

Next generation DNA sequencing is becoming an increasingly important and ubiquitous tool. Sequencing results can be life-changing; they are used to make medical decisions, determine paternity, and are instrumental in forensics. The end-to-end sequencing pipeline, including sequencing protocols, sequencing instruments, and digital analysis, have been be changing rapidly as next-generation DNA sequencers have become exponentially faster and cheaper^13^. However, the risks of adversarial manipulation to sequencing applications are understudied. These innovations and our increasing reliance on DNA sequencing motivates us to study the security of the sequencing pipeline before the technology matures.

DNA sequencers fundamentally are interfaces that bridge molecular information (stored in DNA) and electronic information (encoded digitally). Recent results^14^ demonstrated risks to the digital side of DNA sequencing and subsequent downstream analysis by showing that it was possible to compromise computer systems with malicious DNA strands that encode malware. In this work, we demonstrate that the molecular side of DNA sequencing is also vulnerable to adversarial manipulation. A deliberately crafted DNA sequencing library can be used to modify sequencing results in other samples that are pooled and sequenced together to cause targeted misgenotyping.

Illumina’s sequencing-by-synthesis platforms support multiplex sequencing so separate samples can be sequenced together to increase throughput and scalability. This is done by adding indexes, also known as barcodes, to each DNA fragment during library preparation^2^ The samples are demultiplexed after sequencing by computationally partitioning them based on the sequence of the index. Improper demultiplexing of reads into the wrong bin, known as *index cross-talk* or *index misassignment*, has been a recurring issue since multiplex sequencing was developed^3^ (Fig 1a).

**Figure 1:**
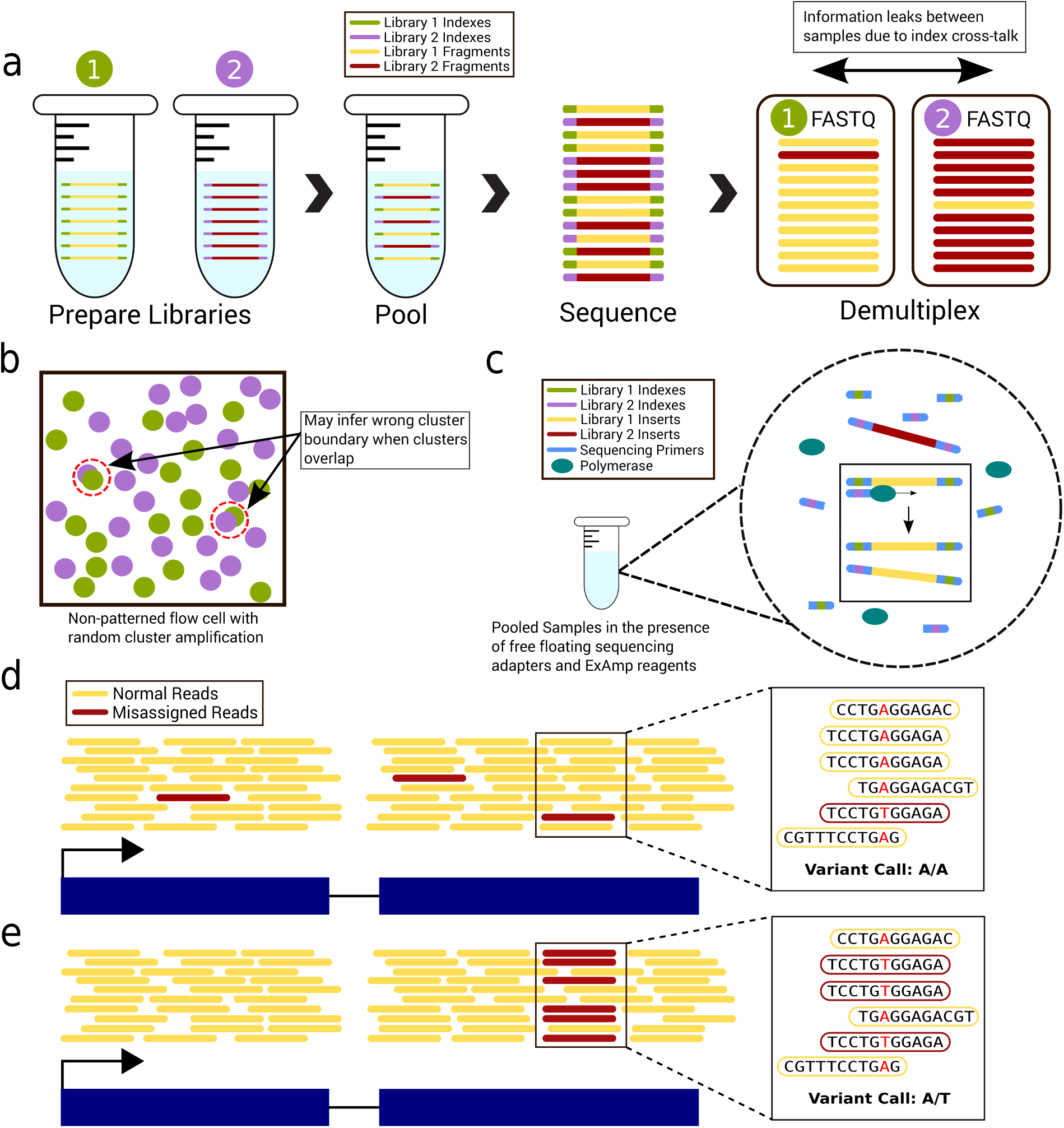
Index cross-talk creates a vulnerable information side-channel. **A**, Index cross-talk between multiplexed libraries leaks information between FASTQ files. **b**, Clonal clusters may overlap on non-patterned flow cells leading to misassignment. **c**, Free floating index primers and polymerases can cause index misassignment. Indexes will prime the 3*′* end of library molecules, which are then extended by DNA polymerase to create mixed index molecules. **d-e**, When misassigned reads are randomly distributed (**d**) they are treated like normal sequencing errors that are unlikely to affect downstream results. However, when misassigned reads are non-uniform (**e**) then they can influence downstream results, in this case, altering a variant call.

Index sequences are designed to be robust to random noise like base substitutions ^15^ and other errors like insertions and deletions^16^. However, using well designed index combinations does not eliminate index cross-talk^5^. It can be introduced by contamination during ultramer synthesis or library preparation and when clusters overlap on flow cells that use random cluster amplification (e.g., MiSeq, NextSeq, HiSeq 2500)^3, 4^ (Fig 1b). Recent issues have been reported with patterned flow cell sequencers that use a new exclusion amplification (ExAmp) cluster generation chemistry^6, 11, 17–22^ (e.g., HiSeqX, HiSeq 4000 and NovaSeq). ExAmp-based cross-talk occurs when residual free indexing primer in the pooled samples primes fragments, which are later extended by ExAmp reagents^6, 11, 17^ (Fig 1c). Costello et al. hypothesize that index cross-talk can occur whenever multiplexed libraries are amplified together due to the presence of residual adapters and polymerases^11^. In response to these issues, Illumina released recommendations to reduce cross-talk that include best practices for the library preparation workflow^17, 23^.

Cross-talk rates vary based on flow cell type (0.2-6.0 % patterned; 0.05-0.2 % nonpatterned) and library preparation method (3-12x higher with PCR-free compared to PCR-plus preps)^5, 11, 17^. Recently, new indexing strategies have been developed that use unique dual indexes to drastically reduce cross-talk rates on patterned flow cells (<0.01 %)^11, 12^. Index cross-talk presents problems for sensitive application that are less robust to errors, like single-cell sequencing^6^. However, it has not been an issue with more robust applications, like high-coverage genotyping, where it is acceptable to use single index demultiplexing^12^.

Index cross-talk causes the inadvertent exchange of information between samples that were intended to remain separate. In the computer security research community, information leakage in unexpected ways is called a *side-channel*^7–10^. Bioinformatics analysis utilities, like variant callers, are somewhat robust to misassigned reads because they are designed to handle random sequencing errors (Fig 1d). However, we hypothesize that sequencing applications, even those robust to index cross-talk, can be manipulated through this side-channel to affect downstream results (Fig 1e). This was suggested as a possibility^14^ but has not been demonstrated until now.

We find that we can make use of index cross-talk with a maliciously designed sequencing library to alter specific variants that are called in other multiplexed samples. This simple attack can be done by sequencing a short amplified fragment that aligns to a single locus; all of the misassigned reads from this sample will align to this same locus in the other multiplexed samples and represent the variant encoded in the sequence of the fragment.

To demonstrate this, we targeted a single-nucleotide polymorphism (SNP) responsible for sickle-cell trait — an A to T substitution in the 6th codon of the *β-globin* gene (dbsnp:rs334). The *β-globin* gene is transcribed in the negative direction, so to be consistent with the variant caller, we subsequently describe this SNP in the positive orientation (i.e., T is wild-type and A is sickle-cell). We made the malicious library from a 400 base pair (bp) synthetic DNA ultramer that was identical to the first exon and promoter of the human reference *β-globin* gene except it contained the sickle-cell SNP. The sickle-cell ultramer was designed to be larger than the typical insert size so there would be many unique inserts after fragmentation. This is necessary because identical fragments would be flagged as PCR duplicates and removed prior to variant calling. The sickle-cell ultramer was prepared into an 8 bp dual-index library, which was pooled and sequenced along with a human exome sample (NA12878) prepared with a single 8 bp index library (current commercial exome library preparation kits support only single indexes). Since NA12878 is homozygous wild type at rs334, a heterozygous or homozygous sickle-cell SNP call would indicate a successful attack (Fig 2a).

**Figure 2:**
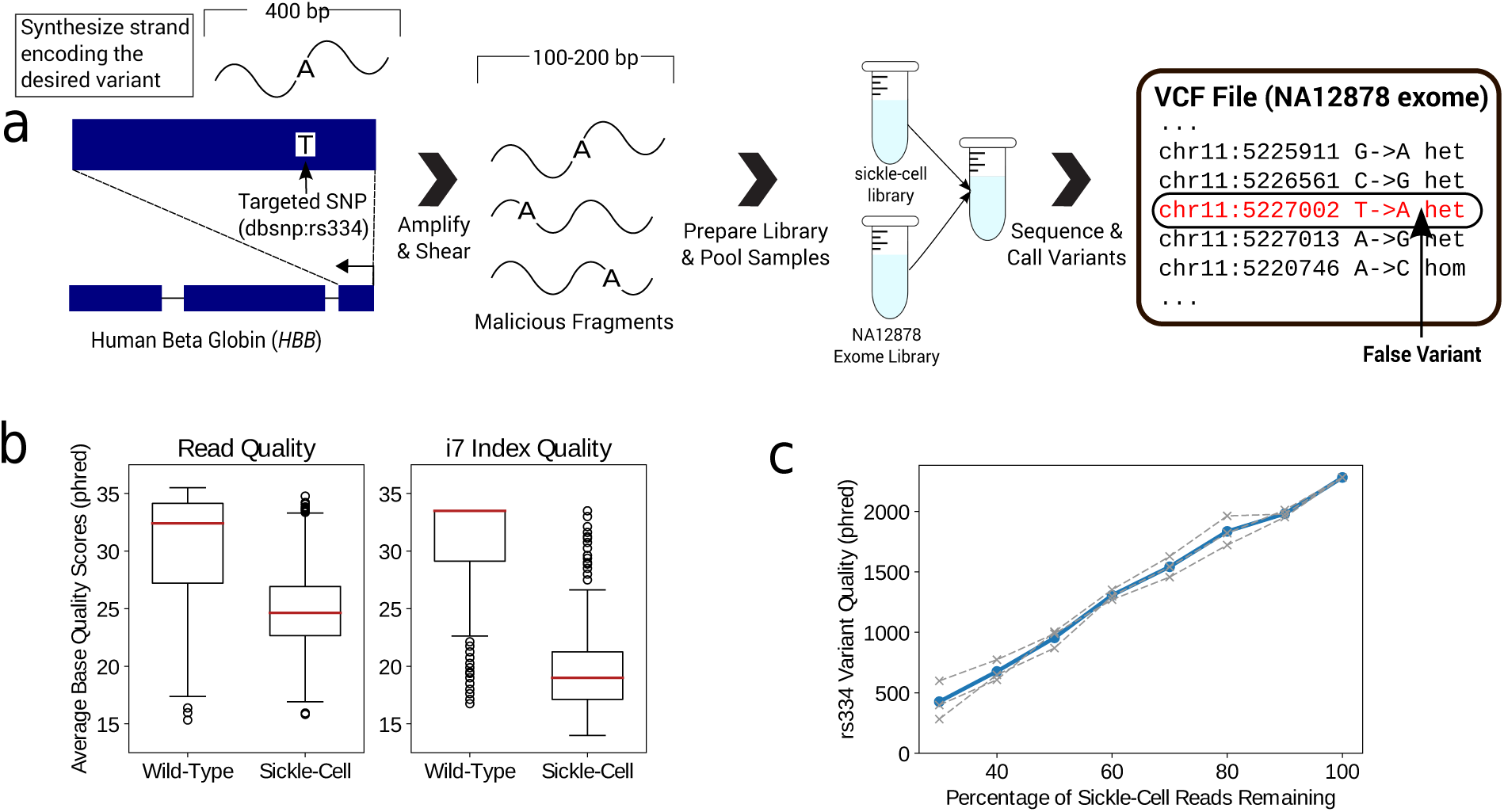
Index cross-talk can be used to modify targeted variants in pooled samples. **A**, To modify a specific variant in another sample, the attacker generates a short ultramer identical to the region of interest, except it contains the desired variant. The ultramer is amplified, fragmented, sequenced together with the target sample. This results in the desired variant being called in another sample. **b**, Average base quality in the read sequence and i7 index of reads that align to the rs334 locus. Reads containing the sickle-cell SNP have lower quality bases than those with the wild-type base. Box-plot elements: center red line is the median, box limits are the upper and lower quartile, whiskers are 1.5x the interquartile range, and points are outliers. **c**, Reads that aligned to the rs334 locus with the sickle-cell SNP were removed randomly in varying proportions to simulate lower levels of misassignment. Each separate simulation is shown by the grey dotted lines; average is shown by the blue line. A heterozygous sickle-cell variant was called as long as at least 30 % of the reads remain. (When 30 % remained, the QualityByDepth*<*2 filter failed in two simulations.)

The two libraries were sequenced in a NextSeq 500 using a Mid-Output flow cell (a non-patterned flow cell with random cluster chemistry). The exome sample was demultiplexed and aligned to the human genome (hg38) using the bwa-mem aligner. SNP and indel variants were called in the NA12878 exome sample using GATK HaplotypeCaller. Index cross-talk from the sickle-cell library led to a false, high-quality heterozygous sickle cell variant call (2281 phred quality score) at the rs334 locus in the NA12878 exome sample. Since only two samples were sequenced, the average coverage was high (321X). The read depth at the rs334 locus was especially high (820), likely the result of cross-talk, since 559 (68 %) of the reads had the sickle-cell base (Supplementary Table 1).

Similar to what has been seen in other work^5, 6, 11^, some reads had invalid dual-index combinations (i.e., i7 index from one sample and i5 index from another). Since the exome library was not prepared with an i5 index, we could only detected mixed indexes where the i7 index comes from the NA12878 exome library and the i5 index comes from the sickle-cell library. A small number reads (758 or 0.0004 %), had this mixed index pair. Of these, 65 % aligned to the 400 bp region used to design the sickle-cell ultramer. The reads containing the sickle-cell base that aligned to the rs334 locus also had significantly lower base quality scores, in both the read sequences and indexes, when compared to reads with the wild-type base (Figure 2b).

To understand variant sensitivity to lower levels of index cross-talk, we randomly removed reads that both aligned to rs334 and had the sickle-cell base. Variants were then called as before. As expected, the quality of the heterozygous sickle-cell variant went down in proportion to the number of sickle-cell reads that were removed (Figure 2c). A false variant passing all quality filters was called as long as at least 40 % of the sickle-cell reads remained, which suggests that the variant would have been modified with substantially less cross-talk than we observed.

Since index cross-talk can be used adversarially, we recommend that unique dual indexes be used whenever pooling independently sourced samples to reduce misassignment, especially in important applications like medicine or forensics. In cases where index cross-talk cannot be eliminated, it is helpful to develop strategies that mitigate or detect when variants have been manipulated. One approach is to filter reads by index quality score. Our results match others^5^ and indicate that misassigned reads have lower index quality scores then normally demultiplexed reads. Therefore, filtering out such reads may remove the false variant without substantially altering true variants. When reads with an average i7-index base quality score of less than 22 were removed, the false sickle-cell variant was no longer called. However, other nearby, downstream SNPs calls were not substantially affected (Fig 3a).

**Figure 3:**
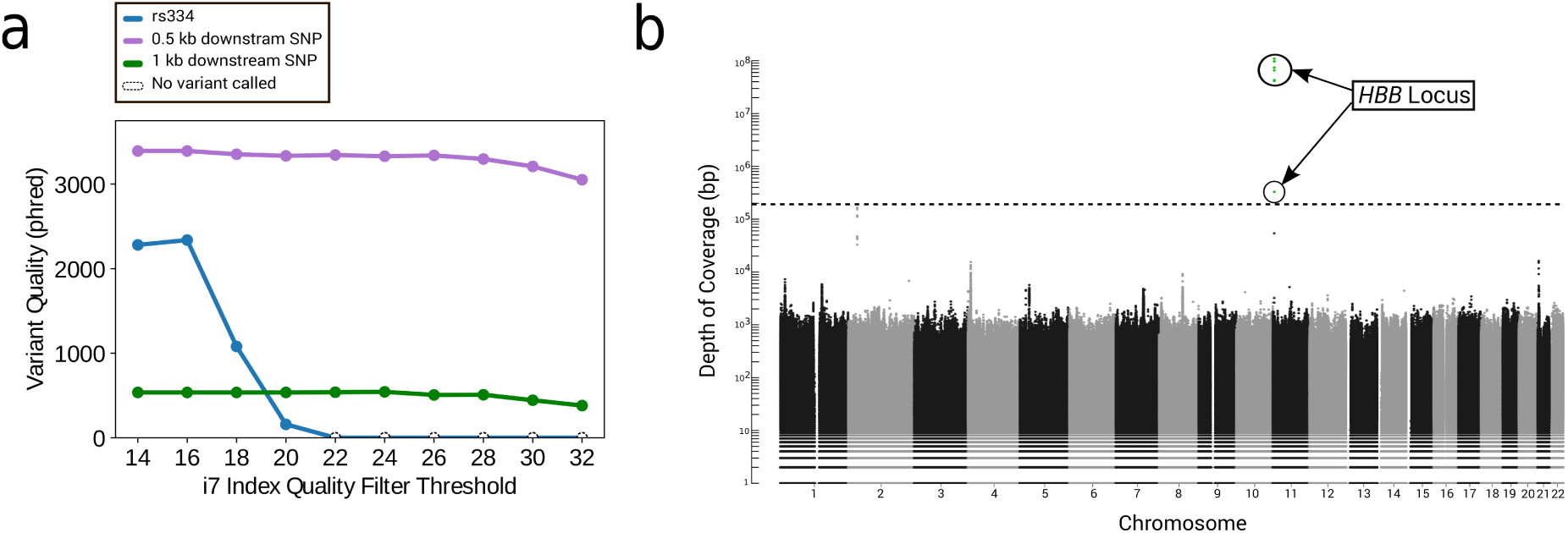
Defenses to prevent variant manipulation. A, Reads were filtered out using the average base quality of the i7 index. Filtering out reads with low index quality scores can be used to remove false variants. By comparison, nearby SNPs (located at *chr11:5226561* and *chr11:5225911*) were largely unaffected by filtering. **b**, Read depth along the autosomal chromosomes using all reads (no demultiplexing). Each point represents the depth at a single position (points with zero-coverage are not displayed). Dotted horizontal line shows the highest coverage not located at the *HBB* locus. Circled green points are in the 400 bp region used to design the sickle-cell ultramer.

This attack requires that the malicious library have highly non-uniform coverage at the desired locus. Therefore, a simple method to detect variant manipulation is to compute coverage at every loci with all reads (no demultiplexing) and look for outliers in coverage. In our sequencing run, the read coverage at the *HBB* locus was over 100 million base pairs, which was 3 orders of magnitude higher than any other position (Fig 3c). Another approach is to call variants using only the reads that demultiplex to mixed-index pairs because those reads represent what will be misassigned into other samples. When we did so, the only variant that was called was a homozygous sickle-cell SNP.

These results show that information side-channel vulnerabilities in high-throughput DNA sequencers can be used to manipulate sequencing results in a targeted manner. Engineers should be aware of side-channel risks whenever combining samples for increased data density and throughput. We encourage the bioinformatics and genomics community to better understand potential adversarial actions against the genomics-information processing pipeline, so we can develop solutions while the technology is young and before problems arise.

## Methods

### Library Preparation

The sickle-cell ultramer and primers for amplification were ordered from IDT (see Supplementary Table 2 for sequence and primers). It amplified with primers, 100 L of 2x Kapa HiFi enzyme mix, 80 L of molecular grade water, 5 L of each primer at 10M diluted in 1x TE buffer, and 10 L of the synthesized ultrame r at 1 ng/L diluted with 1x TE buffer, for a mixture totalling 200 L. The mixture was vortexed on a benchtop vortexer for 10 seconds, then split into two 0.2 mL PCR tubes and placed in the thermocycler with the following protocol: (1) 95C for 3 min, (2) 98C for 20 s, (3) 60C for 20 s, (4) 72C for 30 s, (5) go to step (2) 11 additional times for a total of 12 cycles, and (6) 72C for 30 s. The resulting product had no side products when examined with a QIAGEN QIAxcel fragment analyzer, and it was approximately 165 ng/L.

The human genome NA12878 was ordered through Coriell Institute and was not modified prior to shipping to Genewiz for library preparation.

Both the whole genome and the sickle-cell amplicon were sent to Genewiz for further preparation. The whole genome was prepared with the Agilent SureSelect Exome library preparation kit (v6) to prepare only the exome for sequencing, using index A11 with sequence *CCAGTTCA*. Fragment sizes ranged from 290 bp to 784 bp as measured by the Quiagen Fragment Analyzer. The amplicon was prepared with fragmentation using the NexteraXT kit, using index N703 with sequence *AGGCAGAA* and index S516 with sequence *ACTCTAGG*. Fragment sizes ranged from 168 bp to 608 bp (using the Quiagen Fragment Analyzer).

### Sequencing

The prepared exome and sickle-cell samples were found to be 6.2 ng/L (23 nM) and 2.2 ng/L (11nM), respectively, with the Qubit 3.0 fluorometer. The run was 48 percent exome sample (0.9L) and 48 percent amplicon sample (2L), with a 4 percent PhiX spike-in as a sequencing control. Samples were diluted and denatured prior to sequencing using the NextSeq System Denature and Dilute Libraries Guide. Sequencing was done on the NextSeq 500 and used a 300 cycle Mid kit, with 150 cycles in each read and two 8 bp index reads.

### Downstream Processing

All reads were demultiplexed with the Illumina bcl2fastq conversion software (v2.20.0) using the default configuration (one base pair mismatches was allowed). The create-fastq-for-index-reads flag was used to retrieve index quality scores. The exome sample was demuxed with (i7:CCAGTTCA) and the sickle-cell sample with (i7:AGGCAGAA; i5:ACTCTAGG). The reads with the invalid mixed-index pairs were demuxed with (i7:CCAGTTCA; i5: ACTCTAGG).

To call all variants, reads were aligned to the human genome (GRChg38) using bwa-mem (v0.7.15). PCR and optical duplicates were removed with the Picard MarkDuplicates utility (v2.9.0). Base scores were recalibrated with GATK (v3.7) BaseRecalibrator with the following vcf files from the GATK resource bundle: dbSNP v138, OMNI 2.5, HapMap 3.3, and Mills and 1000G Gold Standard Indels. Variants were called with GATK HaplotypeCaller in discovery mode and SNPs were hard filtered according to GATK’s generic filtering recommendations (QD *<* 2.0, MQ *<* 40.0, FS *>* 60.0, SOR *>* 3.0, MQRankSum *<* −12.5, and ReadPosRankSum *<* −8.0). Exome coverage was computed using the bedtools coverage utility (v2.25.0) with the Agilent SureSelect Exome v6 bed files.

### Variant Analysis

Reads containing the sickle-cell SNP were any that covered the rs334 position (chr11:5227002) after alignment and had the sickle-cell base (T) at that position. All such reads were identified and removed in varying proportions from the demuxed FASTQ file to simulate lower levels of index misassignment. For example, to simulate 90 % levels of misassignment, 10 % of the sickle-cell reads (rounded up) were removed, at random, from the FASTQ file. Then the reads were aligned and variants called on the FASTQ file as usual. Simulation was run every 10 % from 0-100 % three times with a different random seed each time.

To filter out reads based on index quality, the average i7 base phred quality score was computed for each read pair. Any reads which were less than the given quality threshold were removed from the FASTQ file. The remaining reads, which passed the i7 quality threshold, were aligned and had variants called as usual. Variants were called using quality filter thresholds from 14-32 (even only).

The read depth Manhattan plot was generated using the manhattan function from the CRAN qqman package (v0.1.4). The read depth was sampled every 50 bp and plotted; positions with 0 read depth were not plotted.

## Data Availability

The raw BCL sequencing files used to generate the demuxed FASTQ files and the VCF for the NA12878 exome sample are available at Zenodo with the following doi: 10.5281/zenodo.1252436

## Acknowledgements

This research was supported in part by the University of Washington Tech Policy Lab, the Short-Dooley Professorship, and the Torode Family Professorship. We thank Sandy Kaplan, Paul Ney, the Molecular Information Systems Lab, and the Security and Privacy Research Lab for helpful discussions and comments on this paper.

## Author Contributions

P.M.N. led experiment design. L.O. performed experiments. P.M.N. led data analysis and the interpretation of results. P.M.N., L.O., K.K., T.K., and L.C. wrote the paper. T.K. and L.C. supervised the work.

## Competing Interests

The authors declare no competing interests.

## Materials & Correspondence

Correspondence and requests for materials should be addressed to P.M.N.

